# A qualitative model of the rod photoreceptor in natural context

**DOI:** 10.1101/050823

**Authors:** Timothée T. Dubuc, Etienne B. Roesch

## Abstract

Given the intricacies of the retinal neural circuit, which bears a striking resemblance to that of the brain, it is proposed that retinal function goes beyond mere spatiotemporal prefiltering. We hypothesise that aspects related to motion detection and discrimination, anticipation and adaptation to environmental and contextual conditions, which have traditionally been ascribed to the brain, may be supported by neurons in the retina. Such early computations may be dependent on compensative and adaptive mechanisms that stem from qualities intrinsic to the retinal neural circuit and its interaction with the environment (neural transduction time, connectivity patterns, regularities in the input signal, temporal dynamics and light variations).

With a view to investigating the contribution of the photoreceptor population to the processing performed by the retina in natural scotopic conditions, we present a continuous model of the rod photoreceptor. Our model permits the reproduction and exploration of a set of qualitative features displayed in vitro, such as excitation-dependent activation level and time-to-membrane current integration. We captured qualitative aspects of key features selected for their presumed importance in early visual function. Further, we subjected our model to extensive parameter sensitivity analyses, aiming to provide a visual representation of their contribution to the observed qualitative behavior.

**Author Summary:** Primate rod photoreceptor cells constitute the very first processing step in retinal function. This layer influences most of the visual field and presents itself as a tightly packed photosensitive array. Yet, no computational formulation of this neuron is suited for large-scale, time continuous modeling. With a view to studying retinal function in natural contexts, we describe a qualitative, continuous model of the rod. We subject this model to parameter analyses against selected behavioral features. We aim to provide a model that can integrate in further experiments, as well as in large scale network simulation.

## Todo list

### Introduction

The intricacy of neural circuits in the retina suggests that it may support quite complex and powerful computations. It is proposed that retinal function goes beyond mere spatio-temporal prefiltering to include aspects related to motion detection and discrimination, anticipation and adaptation to environmental and contextual conditions, which have traditionally been ascribed to the brain [1]. Such early computations may be dependent on compensative and adaptive mechanisms that stem from qualities intrinsic to the retinal neural circuit and its interaction with the environment; such as neural transduction time, connectivity patterns, regularities in the input signal, temporal dynamics and light variations—which can vary up to 9 log scales. These latter aspects of signal processing, which reflect variations in the input signal, may rely heavily on the base layer of this hierarchical network, the photoreceptors.

Photoreceptors are spikeless neurons that react to light by hyperpolarizing and producing a corresponding electrical flow. They interface with the rest of the network through gap junctions rendering a two-way communication channel. There are at least four types of photoreceptors in the human eye, each of them mostly sensitive to a particular range of electromagnetic radiation [2]. Amongst them, the rod photoreceptors represent about 95% of the photoreceptor population [3, 4] and can detect events as small as a single light quantum [5, 6]—assuming complete dark adaptation, i.e. 40 to 50 min [7])—and convert it into an electrical signal spanning 500 ms.

Phototransduction, which is the process of conversion of light into neural signal, is common to all human photoreceptors. Generally, when a photon of light is absorbed, a cascade of events leads to the hyperpolarization of the cell, the bleaching and ensuing recycling of reacting molecules. In the rod, particularly, the absorption of photons kickstarts a chain of chemical reactions where the photo-activated rhodopsin starts a set of two chemical loops that amplify the initial excitation and control it over time [8]. Each loop relies on self-sustained processes and is able to trigger hundreds and a thousand of reagents, respectively. This complex cascade of reactions supports a global metabolic change from a single local event. The second loop will eventually decrease the medium concentration in cyclic Guanosine Mono-Phosphate (cGMP) [7], driving the gating of Na^+^ and Ca^2+^ ion channels and producing a change in neural current.

During this process, the time scale of the event is fundamentally altered. Photoreceptors present activation and deactivation time that are function of the light stimulation intensity. Unlike other photoreceptors, the rod appears to have a fairly slow time course [9]. For a light event of the microsecond, for instance, it may produce a mirror electric event of the order of the demi-second (5 log scales longer) and up to many seconds for a saturating input (see Figure 5). This phenomenon is heavily studied [7,10–12] and, along with its intrinsic electrical properties, has been the foundation to numerous models representative of subsets of behaviors expressed by the rod and its internal machinery. However, the resulting models, which are generally complex and dependent on an interpretation of the chemistry involved [9,11–13], mainly account for exposures to isolated flashes of light or a part of the activation-recovery process. These models are thus remote from the natural interaction with continuous light. Those limitations make these models unfit to be used as a building block for a time-continuous model of the retina. The retina and photoreceptors have indeed evolved to extract, process and represent information about the natural environment as effectively and as discriminatively as possible [14]. A theoretical understanding of the information processing occurring within the retina thus requires a perspective that will account for the natural interaction with time-continuous stimulation, as present in the interaction with natural scenes.

As such, the retinal neural network and its components may be optimised to work in a continuous time referential, where punctual events are meaningless on their own. With this in mind, we hypothesize that part of the circuitry and accompanying cell behaviors will mainly be understood in a time-continuous context as opposed to a single stimulus event. Some observed behaviors, such as the omitted stimulus-response or diverse adaptation phenomenon, are, in essence, only present as the experiment unfolds over a long enough period of time [15–17]. Other illustrations of long-term phenomena can be found in the adaptive sensitivity of the retina to the contrasts and patterns that shape the stimulating light. When exposed to comparatively higher contrast environments, for instance, the sensitivity of the retina decreases as the kinetic of its response accelerates, i.e. supporting contrast adaptation [18]. This mechanism has been shown to occur in two stages: A fast step (≈ 0.1*sec*) of kinetic acceleration and sensitivity decrease and, thereafter, a longer, slow step (of the order of seconds) during which only the sensitivity is affected and keep on decreasing over time.

Similarly, the retina also appears to slowly modulate its sensitivity to surrounding patterns (pattern adaptation). A classic example may be the effect of prolonged exposure to an environment composed of horizontal bars, which decreases the sensitivity of the retina to horizontal features altogether, in favor of an increased sensitivity to vertical features in the environment [19].

In the present paper, we propose a simple, qualitative model of a rod photoreceptor, as a first step towards a more complete understanding of the retina in the time-continuous domain. We focus our efforts primarily on the reproduction of behavioral characteristics of the rod in action, and the effects of the parameters of our model. In what follows, we first review selected models of the rod, before describing our model. We compare responses obtained with our model to behavioral characteristics of the cell as described in the literature, and propose a thorough parameters sensitivity analysis over selected effects found in the signal, namely rise time, decay time, hard bump, light intensity and duration of responses.

### Electrical properties and existing models

Rods have first been studied through the lens of their electrophysiological and electrical properties, by means of electro-retinography [20]. Naturally, many models have been proposed that review aspects of the electrical flow arising upon stimulation. Such models will preferentially represent these effects as resistor-capacitor (RC) circuits, with sets of resistance and capacitance values representing different morphological systems. Other representations draw inspiration from cable equation theory, and treat the cell as a single, long conductor with variable conductance properties. Those models, even if lacking biological accuracy, have been shown to explain how neural current may emerge and be transmitted throughout the cell, up to the synaptic branching [21, 22]. Similar approaches have been used in the study of synaptic current spreading within the retinal synaptic sheet [23, 24].

A birds-eye view of this range of models will show a certain breadth of complexity, from overlapping mechanisms (e.g. convolution filters followed by RC circuit [25]) that aim to emphasise numerical accuracy, to simple models containing a single RC circuit that reproduces the basic low-pass filtering behavior of the rod in response to light. In contrast to models based solely on RC circuits [22,25,26], the least complex models will typically include approximation of the electrical response to light with equations similar to Hodgin-Huxley [27]. A traditional RC circuit is therefore crafted out of one [28] to two [29] continuous current generators, in series, with a variable resistor [24] and with or without leaking resistance.

A noticeable category of electrical models involve replacing the capacitor within the circuit with an inductor [30–32]–dubbed RL circuits. This technique has been used to study the high-pass filtering capacity that arises when rods are connected in a network; see for instance, models of rods in the toad *Bufo marinus* [30] and the salamander *Ambystoma tigrinum* [31]. Even though these models are based on a time-continuous referential and exhibit many kinds of behavior typical of the rods, they share a critical limitation as they lack multiscale-time signal integration. For a very short light event (<< *τ*), RC models are not able to produce a biologically consistent response. In contrast, RL models will efficiently react to short light impulses but will fail to reproduce sustained stimulation.

Finally, Clark et al. [33] proposed a qualitative model of the rod based on the convolution of two mono-lobed filters, dynamically combined in time. This representation is simple enough to allow formal mathematical manipulation, and representative of some characteristic behaviors of photoreceptors. However, the design is such that the output signal is unsuitable for integration into a network of cells, with a view to studying functional computation.

In response to the above limitations, we aim to produce a model able to account for any type of activation in a time-continuous referential while being modular enough to permit the explicit manipulation of its components without damaging its overall properties. We thus designed a multi-step system: 1) a filter representative of the rod physiology and of rhodopsin light-absorption properties, 2) an ODE echo-loop for the time scale change induced by the phototransduction and 3) a RLC circuit representing electrical events at the cell membrane (See Figure 1).

**Figure 1.**
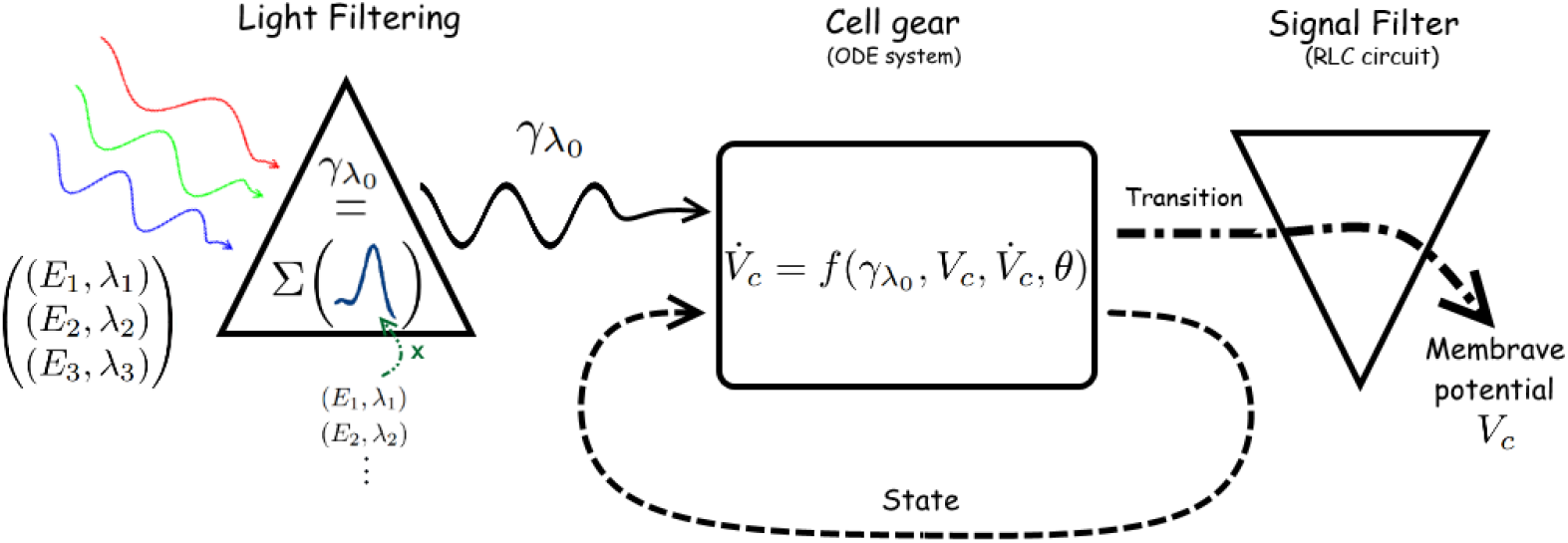
Schematic representation of the rod photoreceptor model. Light is captured through a filter, using Equation 3, which will convert the incoming energy to a number of quantum events *γ* associated with virtual wavelength λ_0_. This quantity represents the energy captured by the rhodopsin for a given incoming light beam. The Cell gear will then use this energy representation along with its current state to compute the membrane potential. The details of this ODE system are discussed in the text.

## Methods

To construct our model, we used data from several sources, referenced where appropriate. When necessary, some components of the model have been optimised using CMA-ES [34] (as implemented in the Python Pybrain library [35]), as this particular algorithm can both be considered as a Monte-Carlo method to avoid local minima problems and an optimization method converging toward a reasonable solution. Our approach is driven by the will to produce a model that permits the exploration of behavioral qualities of rods upon acute as well as sustained light stimulations. We further ambition that our model be useful in the investigation of visual function at both cell and population levels in retinal neural networks. Consequently, we structured our model in such a way as to expose parametric “handles” with enough explanatory power without requiring overly expensive computations. The following gives an overview of the kinds of analyses we are interesting in, and the way we have implemented them.

In addition to constructing a model able to reproduce parts of key behaviors of the rod, we aim to provide a representation of the model parameter space as handles to further behavior alteration (kinetic acceleration, light adaptation, cone model) for analysis. Marder and Taylor proposed a qualitative formalism to quantify the effect of the parameters of a model over a set of monitored effects [36]. For each parameter, they represent a circle, the surface of which corresponds to its impact over the monitored effect. In the present work, we extend this method to represent and explore the neighbourhood of the best known parameter values. We map the surrounding range of values along the horizontal diameter axis of a circle, and locally changed its area in function of the amplitude of the effect studied. The resulting ellipse-like blob represents the effect of the different values for a given parameter, while showing local variations of the neighbouring parameter space (see Figure 2). This method serves in the study of the model, its stability and robustness, and informs the formal analysis of our equations towards improving and building new models. This representation involves three key aspects: the generation of the neighbourhood values by sampling the parameter space, the quantification of the effect variation and value capping.

**Figure 2.**
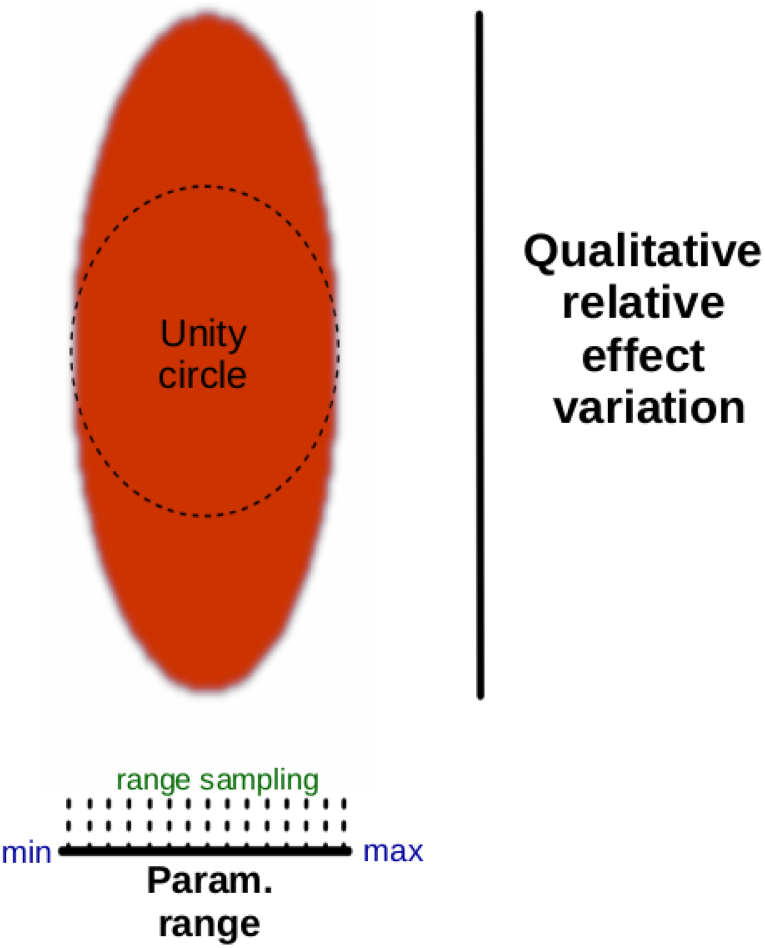
Representation of the relative effect of parameter variation. The parameter variation unfolds along the horizontal axis of the Unity circle 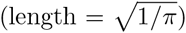. According to the position on this axis, the vertical amplitude of the colored surface represents a measure of the relative effect (See Equations 1 and 2). The unity circle represents a circle with an area equals to 1, and serves as a baseline for interpretation.

### Sampling of the parameter space

We generate a range automatically for a given parameter, in a two-step process. First, the log scale (*ls*) of the optimal parameter value (*opv*) is computed and used to generate a base number (*bs* = 10^*ls*–1^). This number reflects the best known value for that parameter in representing the data. A series of parameter values is then generated by sampling a range evenly, on both sides of the parameter space [*opv* − *bs*/2; *opv* + *bs*/2].

### Effects comparison

The relative influence of each parameter variation over the observed effect has to be measured in a way ensuring numerical and representation stability, as both parameter value and observed effect can yield very large variations. The logarithm of the ratio of the effect variation ranges over the parameter variation range as been chosen as the most stable (low numerical change even in case of large effect fluctuation, no numerical instability as *range_param_*¬0) and most informative to display this relation.

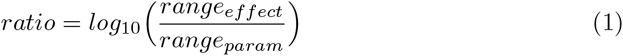

### Capping

For some variables, a small change in parameter value (≈ 10^−4^) can lead to very big fluctuations in term of measured effect (≈ 10^3^) and conversely. Our mapping had to be able to legibly represent all those variations. To achieve this, the log ratio computed in Equation 1, representing the local area of the circle, has to be filtered and capped between two positives values. For clarity, we chose this filter so that a linear dependence between parameter value and effect would be represented as an ellipse, reflecting a one-to-one dependence with a circle of area 1:

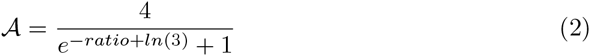

## Our model

We propose a three-tier model that decomposes important functional aspects of rod in human (Figure 1). Schematically, the light is first incorporated to the model through a mathematical filter, representing rhodopsin absorption. Second, relying on differential equations, we designed an echo-loop aimed to change the time scale of the initial data flow accounting for natural chemical inertia. This echo loop finally drives a gating resistance in a RLC circuit standing for the neuron membrane. This allows us to replicate a wide set of natural behavior, whilst running simulations at different time resolution—see complete list of parameters in Table 1.

### Physiological properties of phototransduction: Light filtering

The first component is a filter representing the rod physiological properties that convert the different incoming light radiations (*γ_i_* photons of wavelength *λ_i_*) to a virtual equivalent quantity of energy λ_0_. To create this filter, we extracted parameters from the original data by Wald and Brown [10], representing rhodopsin photoactivation rates, in the form of a mixture of exponential functions (see Figure 3):

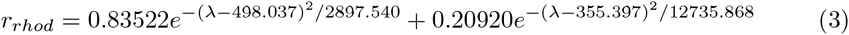

**Figure 3.**
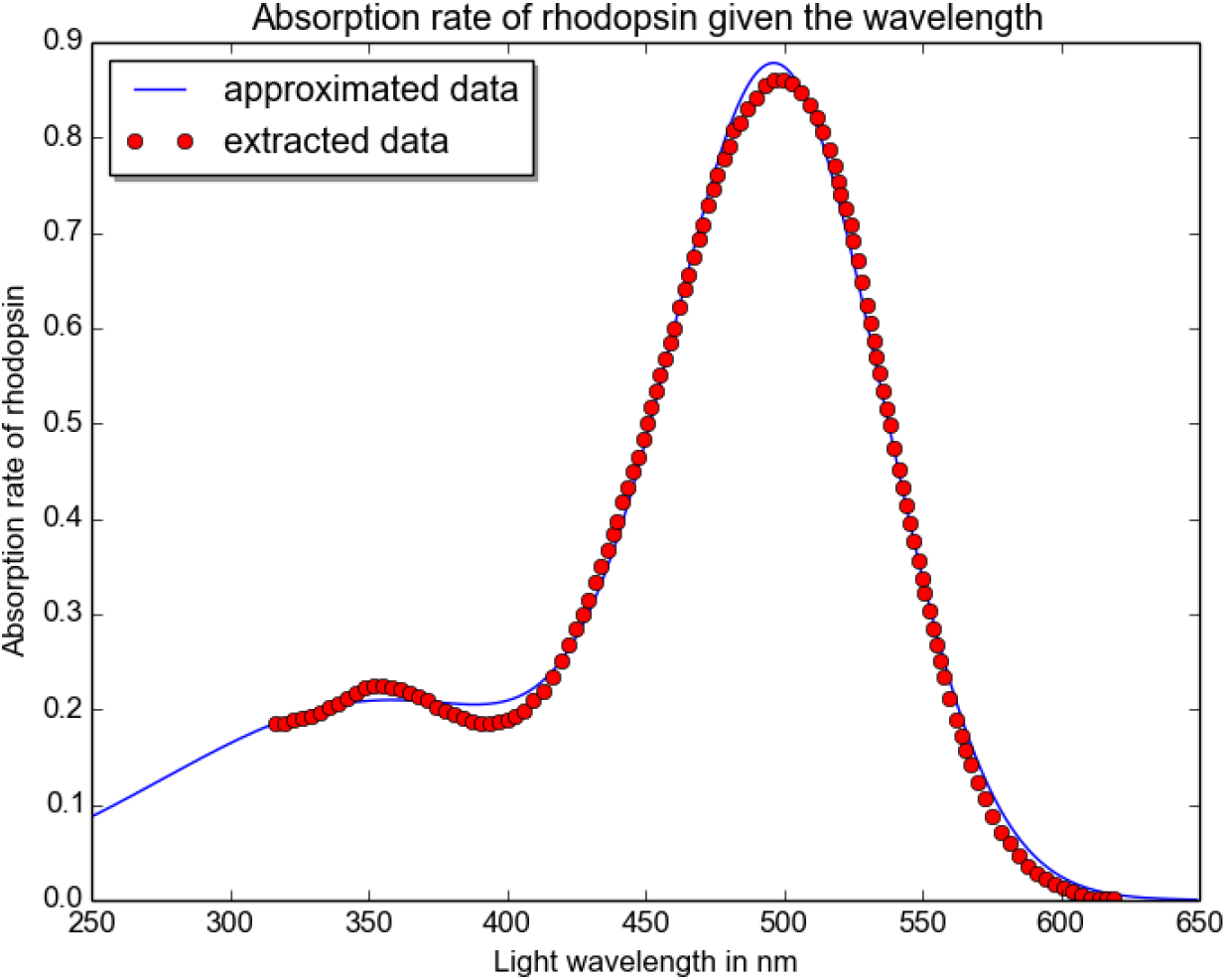
Absorption rate of the rhodopsin given the wavelength. On this graph, the red dots represent the data extracted from the graph in [10]; and the blue line the extrapolated continuous function. As visible, the learned function does not exactly match the experimental data: the qualitative nature or our work disregards high numerical accuraty in favor of computation simplicity.

The quantity *r_rhod_* is then used to produce a representation *γ*_λ_0__ of the global number of photon absorbed by the rhodopsin as function of the dimensions for a given cell [37]:

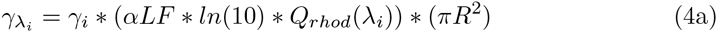

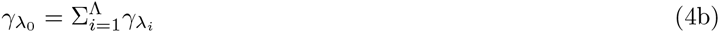

With *α* the specific axial pigment density of the rhodopsin (0.016 *μm*^−1^ in primates [38, 39]), *L* the length of the rod outer segment (25 *μm* in primates [39]), F the absorption ratio of the light due its polarization, *ln*(10) the naperian extinction coefficient [40] and *R* the radius of the rod (1*μm* in primates [39]). F is equal to 1 in case of polarized light in the optimal direction, but is here 0.5 to account for unpolarized light.

*Q_rhod_*(*λ_i_*) represents the quantum efficiency of the rhodopsin (with *Q_rhod_*(*λ_i_*) = *r_rhod_*(*λ_i_*) * *q*) and *q* the quantum efficiency. This value has been determined to be wavelength-, concentration-, and temperature-independent and included within the interval [0.6; 1 [. *Q_rhod_*(*λ_i_*) has been found to be 0.67 (as per [41, 42]) for a 500λ light exposure; For *r_rhod_*(500) ≈ 0.875, we chose *q* = 0.766).

The final value *γ*_λ_0__ gives the raw number of photon absorbed by the rhodopsin and hence the number of isomerisation igniting the phototransduction process within the cell.

### From light to signal: Cell gear

The second component is responsible both for the driving force at the level of the cellular membrane, and the change of time scale that occurs in the system, from microseconds to second-long signal. The reproduction of this last characteristic is important, as we aim to produce a model in the time-continuous domain, which is able to respond to a constant flow of input.

Inspired by the FitzHugh-Nagumo model [43], in the way that it steers the entire system by leaning on a curve (here a hyperbole), we introduce a scale change through an ODE loop with positive feedback acting as an echo chamber. The incoming light stimulus *γ*_*λ*_o__ is first cast to another space (Eq 5a). This casting ensures that a photon quantity in the interval [0,1[ gives a negligible amount of response while producing a tonic monotonous capped response on 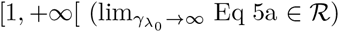 ensure a stable reliable input signal. The resulting quantity is then injected in the loop (Eq 5b and 5c). The limits and general behavior of the system are defined by the equation 5c. Equation 5b drives the total duration of the echo signal from a micro/millisecond excitation to a second(s) long response. We engineered this echo-loop so as to reproduce a set of qualitative features:

- The system must react to a single flash of light by describing a complete hysteresis trajectory.
- The system must display a constant, non null, proportional activation to a steady light exposure.
- The system must reset to its resting point (*a* = 0 and *v* = 0) if not excited.

The resulting equation is composed of three sub-components that ensure the stability of the system: The first sub-component (excitation) provides the system with a fast reaction following a light impulse (*h*^2^). The second sub-component (normalization) damps the activity to ensure that the overall system will produce the desired hysteresis and equilibrium point when stimulated by a steady light. This is achieved through the term 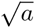 that dampens the response created by the excitation, to then drive the system towards a steady state. Finally, the third sub-component (inhibition) acts as a refractory force that will eventually reset the whole system after a punctual activity. Equation 5c expresses a parabola over which the system activation pans out. Equilibria points are created either at the intersection (*v, a*) = (0, 0) during an absence of light or on the parabola described by Equation 5c for a steady light exposure.

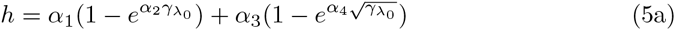

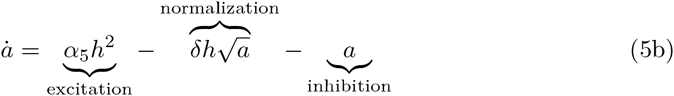

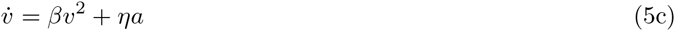

This ODE system generates a trajectory function of the initial energy discharge injected through *γ*_λ_0__: It becomes wider as the intensity of the light grows. The trajectory is also designed to be slightly wider as the lighting time increases for a given light energy. We review the overall effects of these different parameters later in the present paper. The value of *v* will eventually, drive rheostat *R_s_* in ways similar to the effects of cGPM on the ion channel gating in vivo.

### Shape and time fine-tuning: Signal filter to membrane potential

The last component of the model converts the driving impulse given by the ODE loop (see equation 5) to a membrane polarization (or hyperpolarization).

Based upon the idea that the membrane of a neuron behaves similarly as a capacitor, this task is achieved through an RLC-circuit with adjustable gating and leaking resistors (see figure 4). This module will essentially produce a quantity *V_c_*, the rod membrane potential, after the variable *v* expressed in equation 5c.

**Figure 4.**
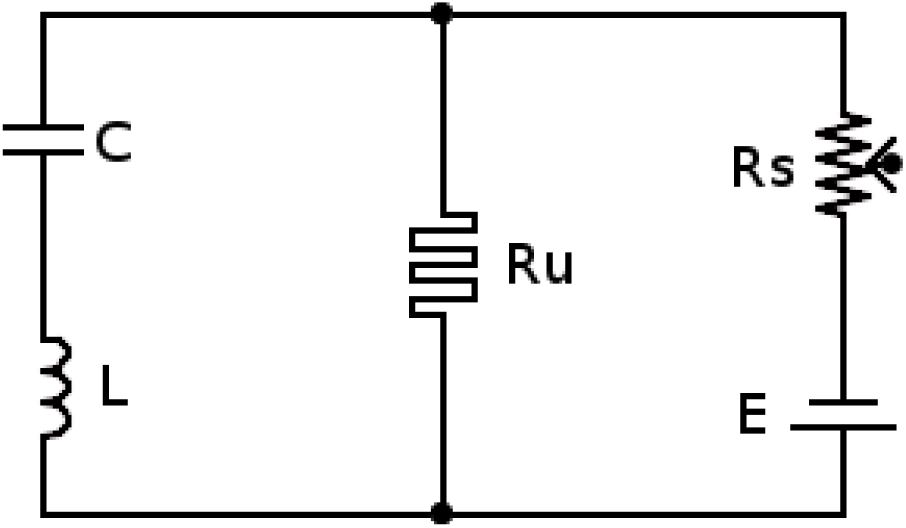
RLC filter block diagram. Circuit representing the membrane part of the model. The potential of the capacitor C represents the rod membrane potential, the leaking memristor *R_u_* (see Equation 8) and the inductor L its inertia, *R_s_* is a switch like rheostat which value drops when exposed to light (see Eq 7).

The mathematical expression of the membrane potential *V_c_* is given by:

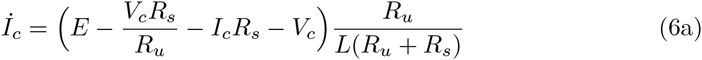

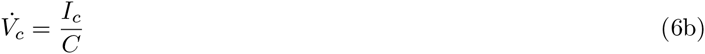

When hit by light, the conductance of *R_s_* increases (see Equation 7) allowing the current to flow inside the circuit and to charge the capacitor.

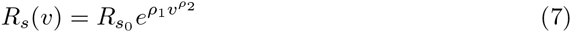

In absence of light stimulus, the value of the resistance *R_u_* increases to account for the longer recovery time displayed by the rod when exposed to high light intensities. This fluctuation, which is similar to a memristor, is given by equation 8

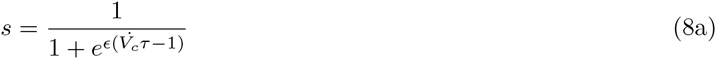

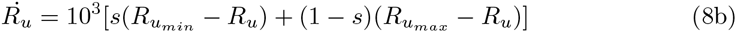

Where *τ* is the time step of the simulation expressed in seconds (here, 10^−6^ *sec*). In accordance to the hypothesis put forth by Rieke and Baylor [5], the resulting *V_c_* signal is clipped (*V_c_* > –1) and normalized, to simulate saturation levels of the whole system.

The resulting overall behavior displays features comparable to a real rod photoreceptor, albeit not numerically accurately (see Figure 4b). From the literature, we identified a number of behavioral features, which may be desirable in such a model, which aims to explore functional effects of neural networks of the retina. In the next section, we review these behaviors, and describe the parameter sensitivity analysis we used to evaluate our model.

### Behavioral similarities with rod cell

The rod photoreceptor is stage for a series of complex chemical mechanisms leading to key characteristics that we are trying to model. For instance, the stronger the light excitation, the faster the reaction and the slower the recovery; or, higher and longer levels of activations leads to more intense responses. For our model, we selected a set of parameters capable of qualitatively reproducing those behaviors (see Table 1), which we take as initial conditions for our parameter sensitivity analysis.

Better results could be achieved with numerical optimization algorithms over the entire parameter set. Unfortunately, this implies a large enough set of data (>> *nbr_parameters_*) to be to able avoid common problems such as over-fitting or the destruction of stability and behaviors. The amount of data available, however, gathered from the literature, is not suited for a black box learning algorithm to perform appropriately. When applied on the entire ODE loop, the stability of the system over long periods of activation appeared to be highly compromised.

**Table 1.**
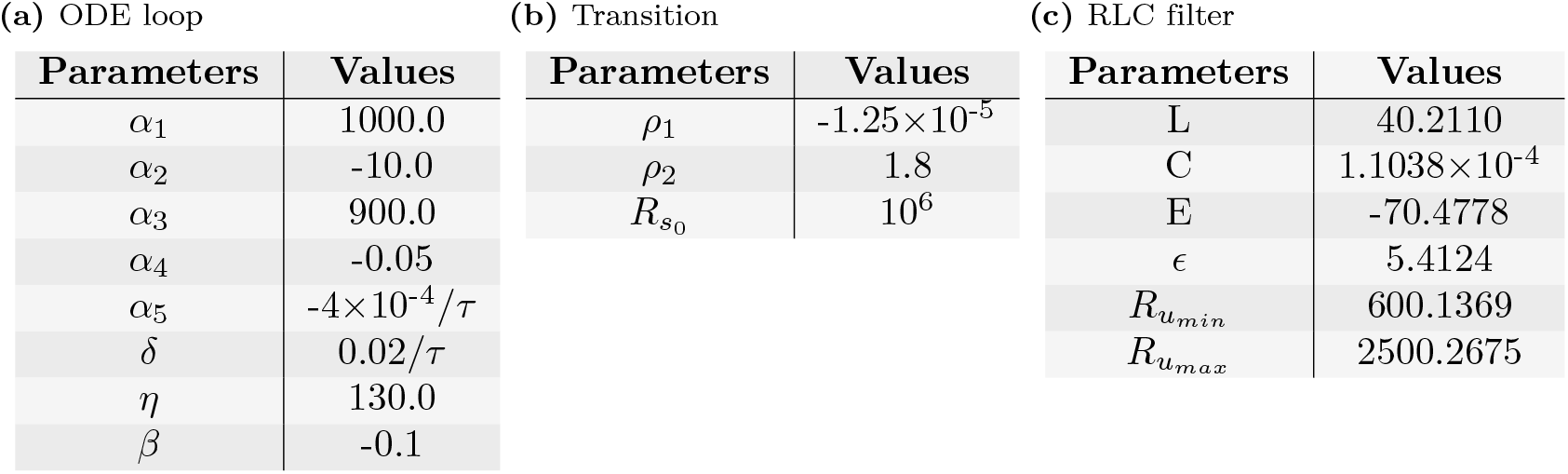
Parameter values. Table showing the selected parameter values for the different parts of the model. Subtable (a) refers to the equation 5, subtable (b) to the equation 7 and subtable (c) to the equation 6 and 8. Note that the results obtained by numerical optimization (Subtable (c)) have only been allowed 4 decimal numbers. *τ* is here the time constant of the model expressed in seconds.

#### Activation and refractory speed

Repeatedly observed in empirical setups, the rod cell response to light appears to have a faster rising curve as the intensity of the light grows. An opposite phenomenon can be observed during the decay time, which becomes longer as the intensity of the stimulation increases. Figure 5 shows the reaction of both a real rat rod recorded in vitro and our model to short stimuli (2 *μsec* flashes) at four different light intensities (1.7*γ*, 29*γ*, 300*γ* and 4000*γ*).

**Figure 5.**
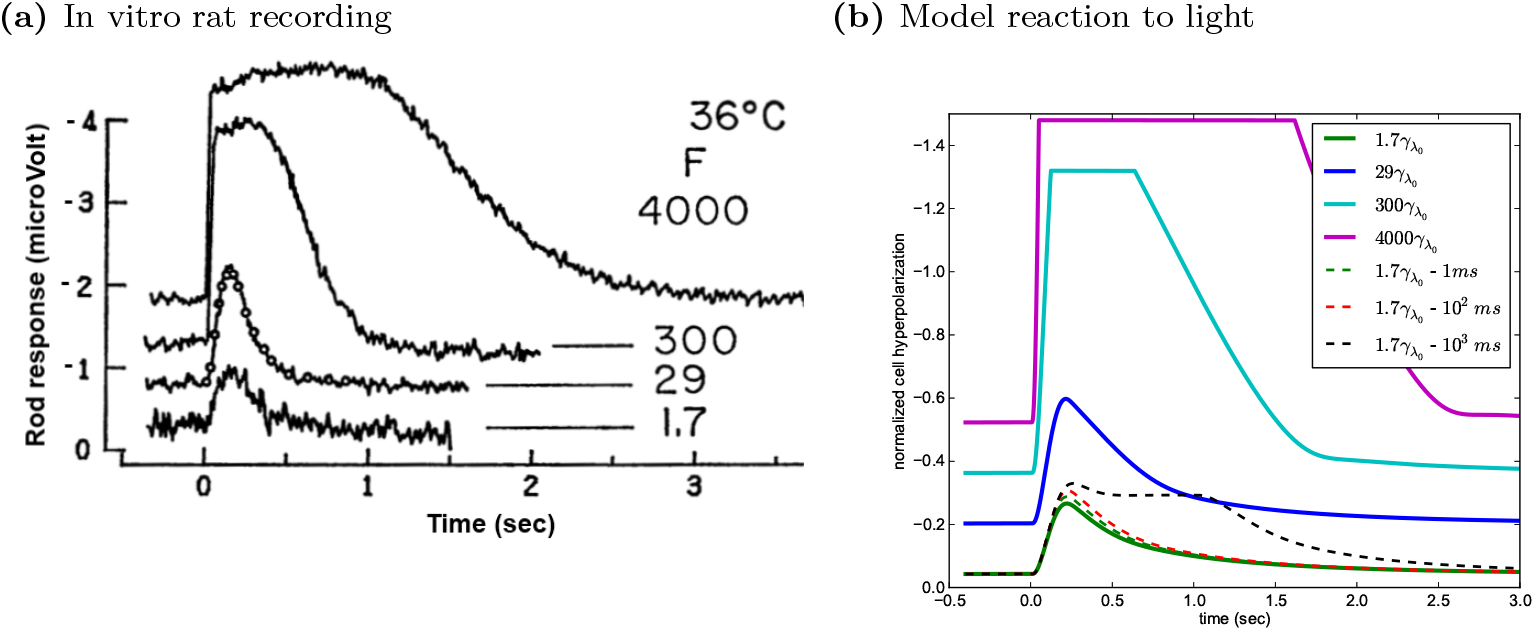
Rod and model reaction to brief light flashes. Comparison of both a real rod and our model reaction to short light stimuli. In both cases, the activation levels before the 0 point of the abscissa axis (flash income) represents the resting potential labelled as 0. (a): Four responses of rat photoreceptor to 2 *μsec* flashes at 36°C in vitro. The intensity of the flash F is expressed as the number of photon absorbed per rod (1.7*γ*, 29*γ*, 300*γ* and 4000*γ*). The circle plots on the *F* = 29 γ curve are produced through Fuortes and Hodgkin’s [26] model and perfectly follow the low light response. (Figure modified from [22]). (b): Output responses produced by our model given the same number of absorbed photon. As visible here, the activation curves present similar characteristics in activation levels, duration and - for small light intensities - shapes.

Those two phenomena seem to suggest an emphasis on the high intensity events, as if labelled as potentially interesting for the visual system. When a high response intensity arises, it tends to mask the low intensity event for a period of time function of the causal light excitation. It is important to note that this temporary damping is not total, as the rod displays a multiple phase behavior that enables it to sustain a small activation for new information and to process it in background until the current level of overall activation finds itself low enough to display them (See section Phase behavior).

#### Level of activation

Another important aspect of the rod response, the reaction level, appears to depend both on the intensity and duration of the incoming signal. Each of these factors seems to have its own degree of saturation. An activation level induced by a long stimulus can find its steady state higher than the response maximum response to a short stimulus (See Figure 7 - last 200 ms). Note that the opposite is not true.

In an attempt to reproduce this behavioral feature, we designed the ODE amplifying loop in such a way that the system is both sensitive to time and to intensity (see Equation 5). For a fixed intensity, the system will create a manifold of trajectories (See Figure 6 - dashed lines) within the range of reaction of the initial intensity based system (Figure 6 - plain lines) but with a slightly different dynamic accounting for the difference in the activation duration. Here, the difference in dynamic is expressed in the shape of the activation plateau within the state space of the ODE loop: the dynamical system phase state is flatter and becomes closer from being perpendicular to the ordinate axis.

**Figure 6.**
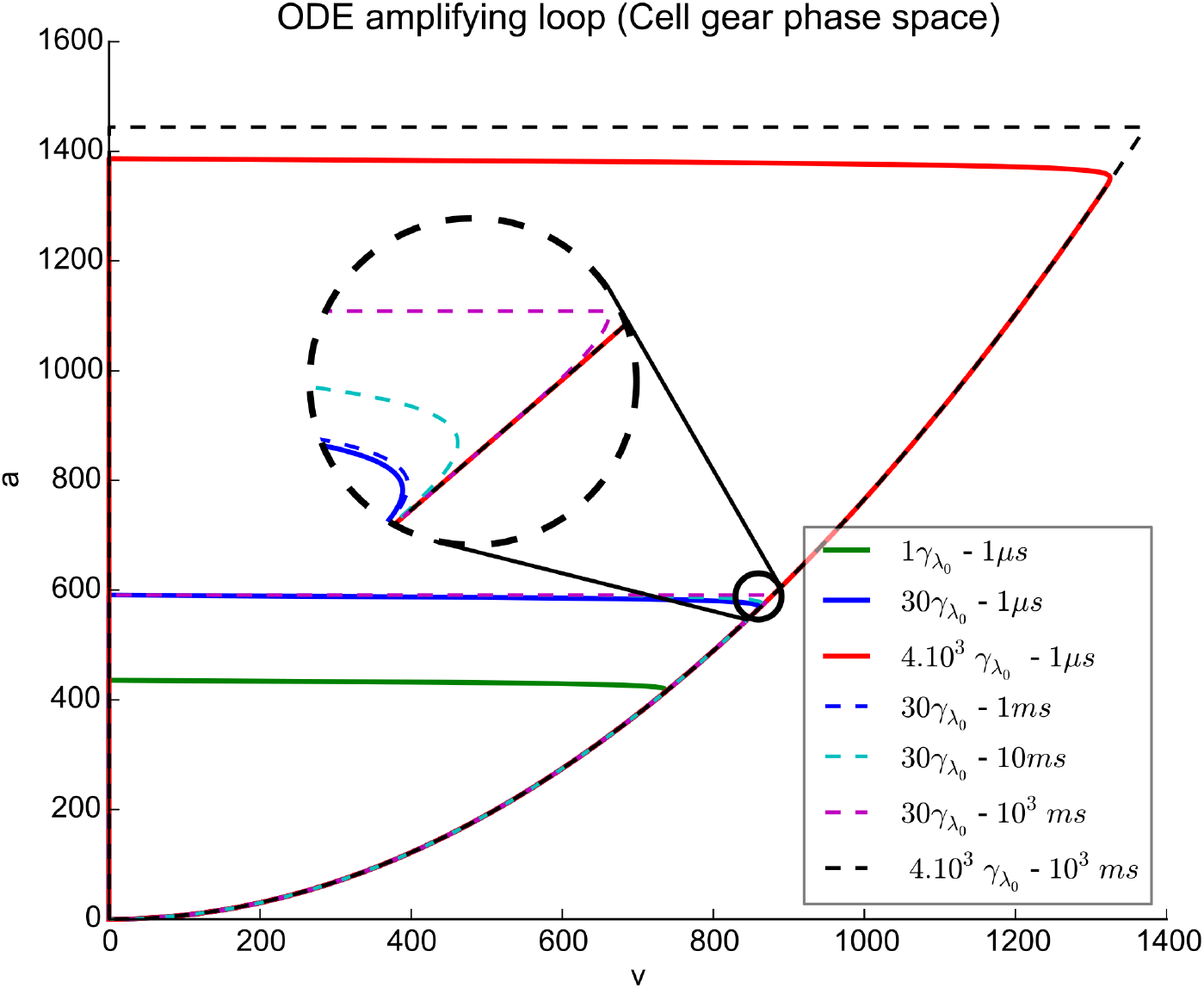
State space of the ODE amplifying loop. Representation of the phase state of the ODE amplifying loop when exposed to light flashes of different intensities for a fixed duration (plain lines) and different durations for a fixed intensity (dashed line). We can distinguish two separated parts: the rising phase whenever *v* increases (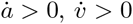 and following the parabola trajectory) and the recovery phase whenever *v* decreases 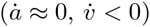. The system equilibrium in case of steady light stimulation appears at the separation point of the trajectory from the parabola induced by the Equation 5c. The phase state displays clear wider refractory orbits as the intensity of the light grows until it reaches a saturating level, here in red, defining the absolute range of possible reaction towards light intensities. However, for each light exposure, a new orbital manifold is created by the model as the excitation duration extends until the plateau of the rise phase (aka: a neat diverging angle from the parabola curve as opposed to a smooth separation curve) becomes parallel to the abscissa axis (magenta and black dashed lines).

#### Constant light response

We aim our model to be sensitive to time-continuous, steady light stimulation. Like purely electrical models, our membrane representation is capable of such a behavior. However, the ODE system has to deliver a constant driving force for the membrane to mirror the steady excitation. This is achieved through the couple excitation-normalization terms of Equation 5b that stands for the equilibrium found in the rod photoreceptor between rhodopsin photoactivation and its deactivation through rhodopsin kinase and arrestin [11]. This same ODE loop (Equation 5) ensures a slow increase of the response as the exposure time grows, until an equilibrium point is reached [37] (See Figure 4b - dashed lines). For long and high enough steady light inputs, in vitro recordings of the rod photoreceptor reveal an initial bump preceding a fixed level activation [37, 44](see Figure 7). Finely tuned capacitance and inductance values enable our membrane representation to display such a feature (see Figure 4b - dashed black line).

**Figure 7.**
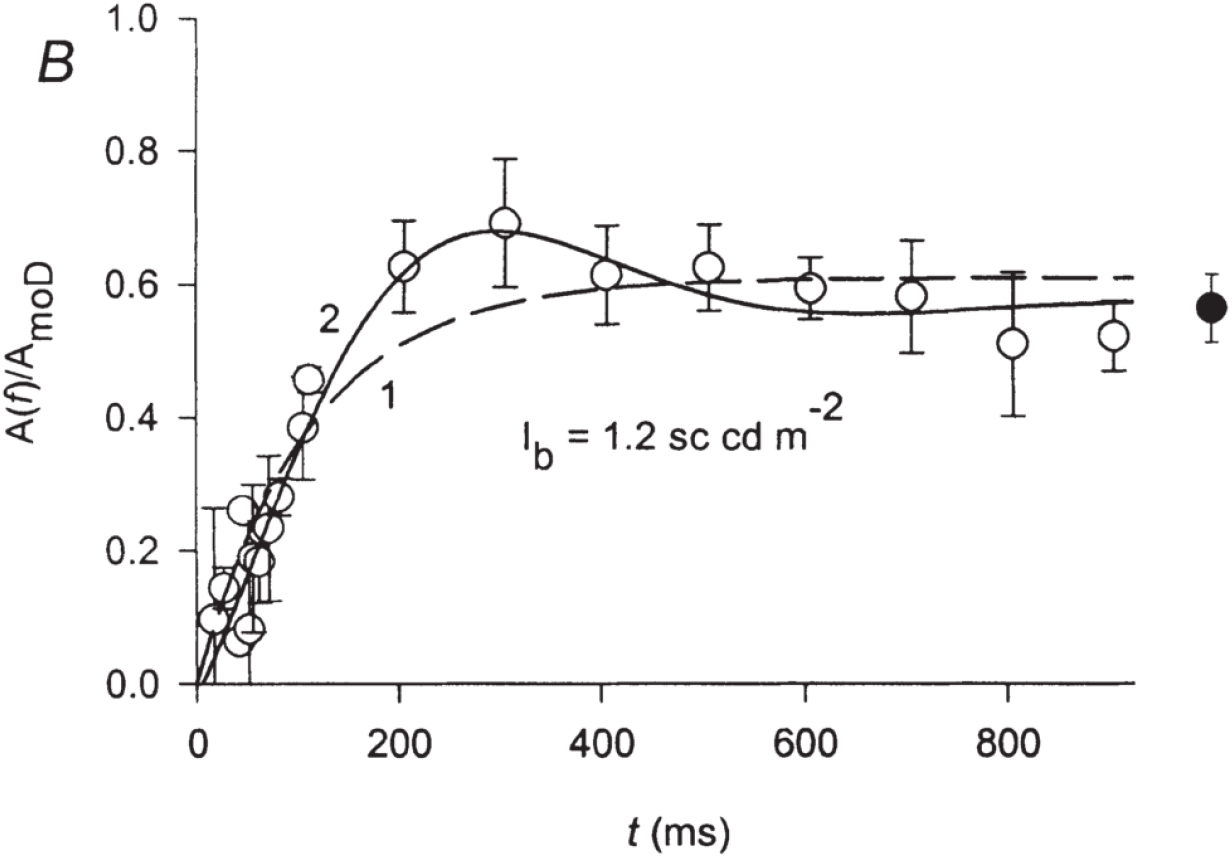
In vitro reaction to steady light. In vitro normalized recording of a mouse rod photoreceptor reaction to a steady light illumination eliciting half steady saturating response (1.2 *sc cd m*^−2^). Empty circles represent the average rod response and standard deviation (vertical bars) over 6 recording. Black full curve is the result of the author’s model. The black circle represent the final activation state achieved by the cell after 2 mn of exposure (extracted from Silva & al. [44]).

### Parameter space

A functional model over a set of parameters is not enough. Having the ability to tweak those parameters and to make the model behave differentially can be as interesting as having an accurate representation of the studied system. Inspired by Marder & Taylor [36], who advocate the need to go beyond the single study of a generic behavior, we monitored a set of behaviors characterizing the rod reaction to light after a brief or constant illumination: Rise time, Decay time, Hard Bump (see Figure 8 for a graphical representation of the monitored behaviours, and respectively Figures 9, 10 and 11 for the results. See Section *Result representation for further details concerning the chosen visual presentation* and Figure 2 for explanations of the data presentation method), Duration-dependent response and Excitation-dependent response (respectively Figures 12 and 13).

**Figure 8.**
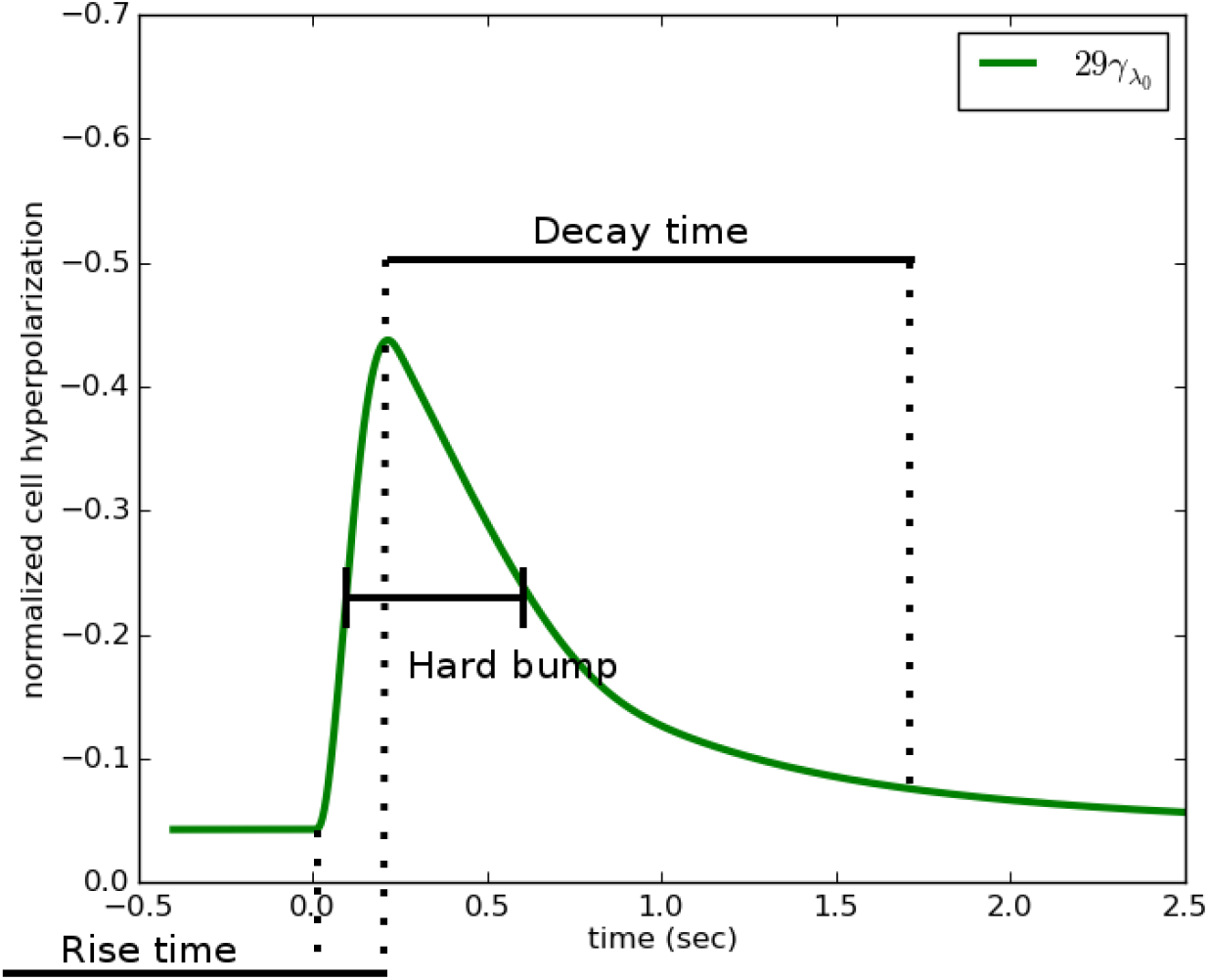
Visualisation of the Rise time, Hard bump and Decay time. Visualisation of three of the five qualitative behaviours studied in this work. The Rise time is the time it takes for our model to reach full hyper-polarisation consecutively to a flash of light, the Hard-bump is the time needed by the model to go from half its rising curve (here *t* ≈ 1000ms) to the same membrane potential on its recovery phase (here *t ≈* 6500ms). Finally the Decay time denotes the entirety of the time needed to move from the apex of the reaction curve to the resting state. The curve here has been extracted from Figure 5 and subjected to a different scaling for the sake of visibility.

**Figure 9.**
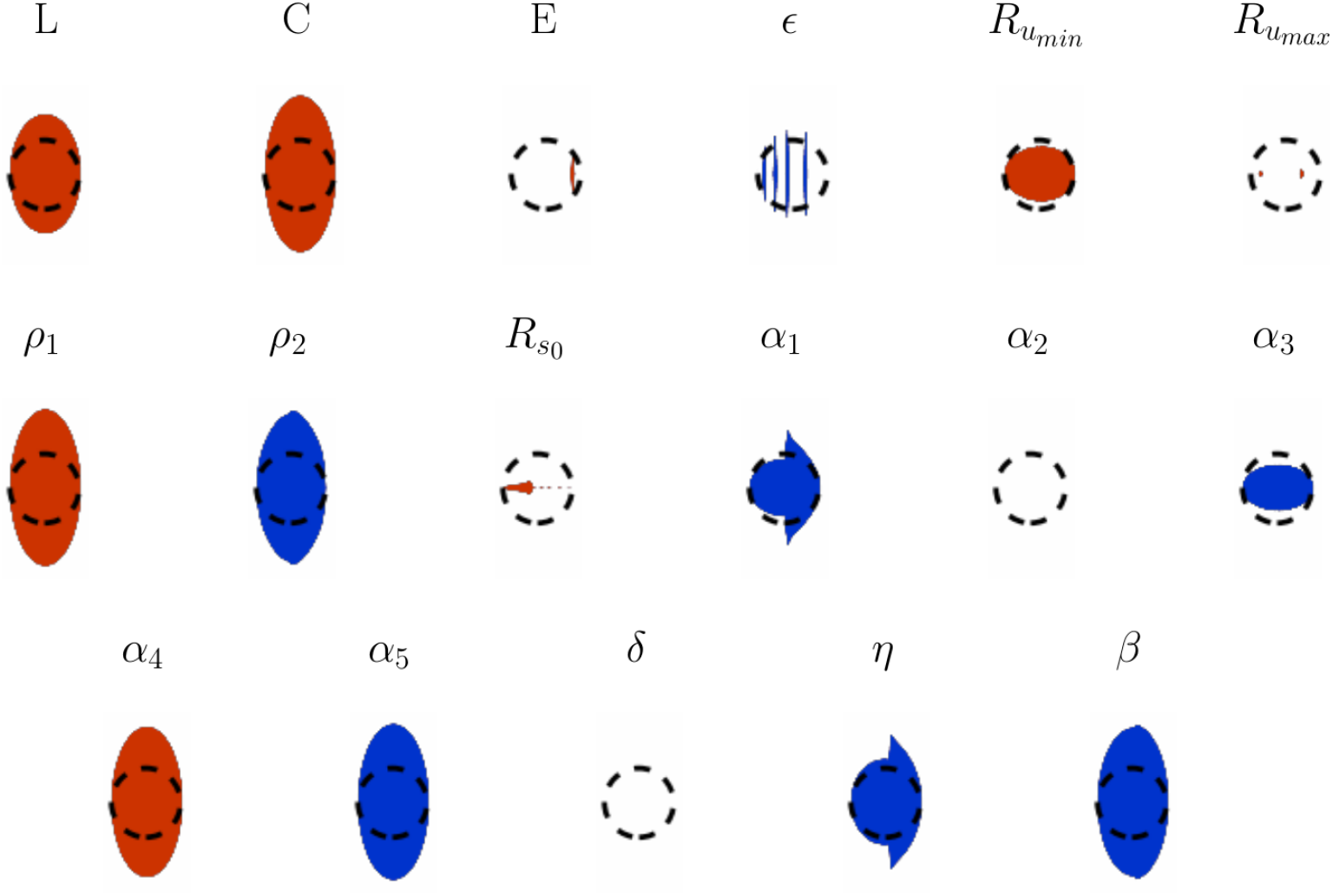
Parameters variation effect over the rise time. For each simulation, the rod has been exposed to a single *1μsec* flash causing the equivalent of 150 rhodopsin photoactivation. This diagram displays 4 non-impacting (*E, α*_2_, *δ* and *Ru_max_*), 2 low impacting (*ϵ* and *Rs*_o_ - the red mass on the right of the blob denotes a very small yet present impact) and 11 strongly impacting parameters. We can notice that among this last group only three of the parameters display a highly non linear parameter-value to effect-intensity relation (*α*_1_, *α*_3_ and *η*) suggesting a plain regular parameter space.

**Figure 10.**
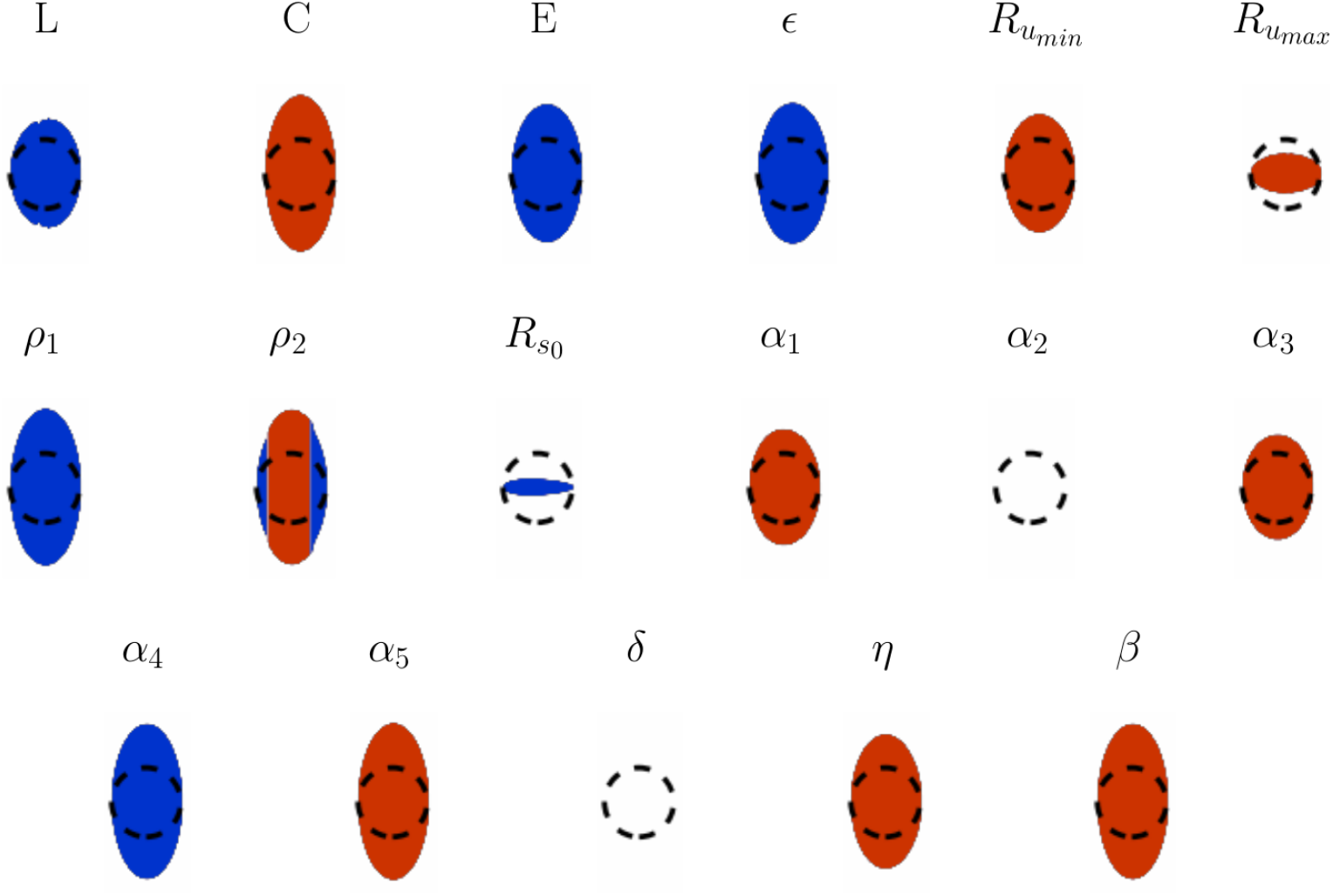
Parameters variation effect over the decay time.

For each simulation, the rod has been exposed to a single 1*μsec* flash causing the equivalent of 150 rhodopsin photoactivation. This diagram displays 2 non-impacting (*α*_2_ and *δ*), two low impacting (*Ru_max_* and *R*_*s*0_) and 13 strongly impacting parameters. Two parameters (*L* and *C*) display non-linear and non-regular parameter-value to effect-intensity. As expected, the inductance value is here at stake. When modified, it can induce current oscillations through the RLC filter circuit and therefore slightly change the moment the algorithm detects the end of the recovery time: even if not stabilized, the voltage derivative used to detect the momentum of the current will appear null by numerical approximation.

**Figure 11.**
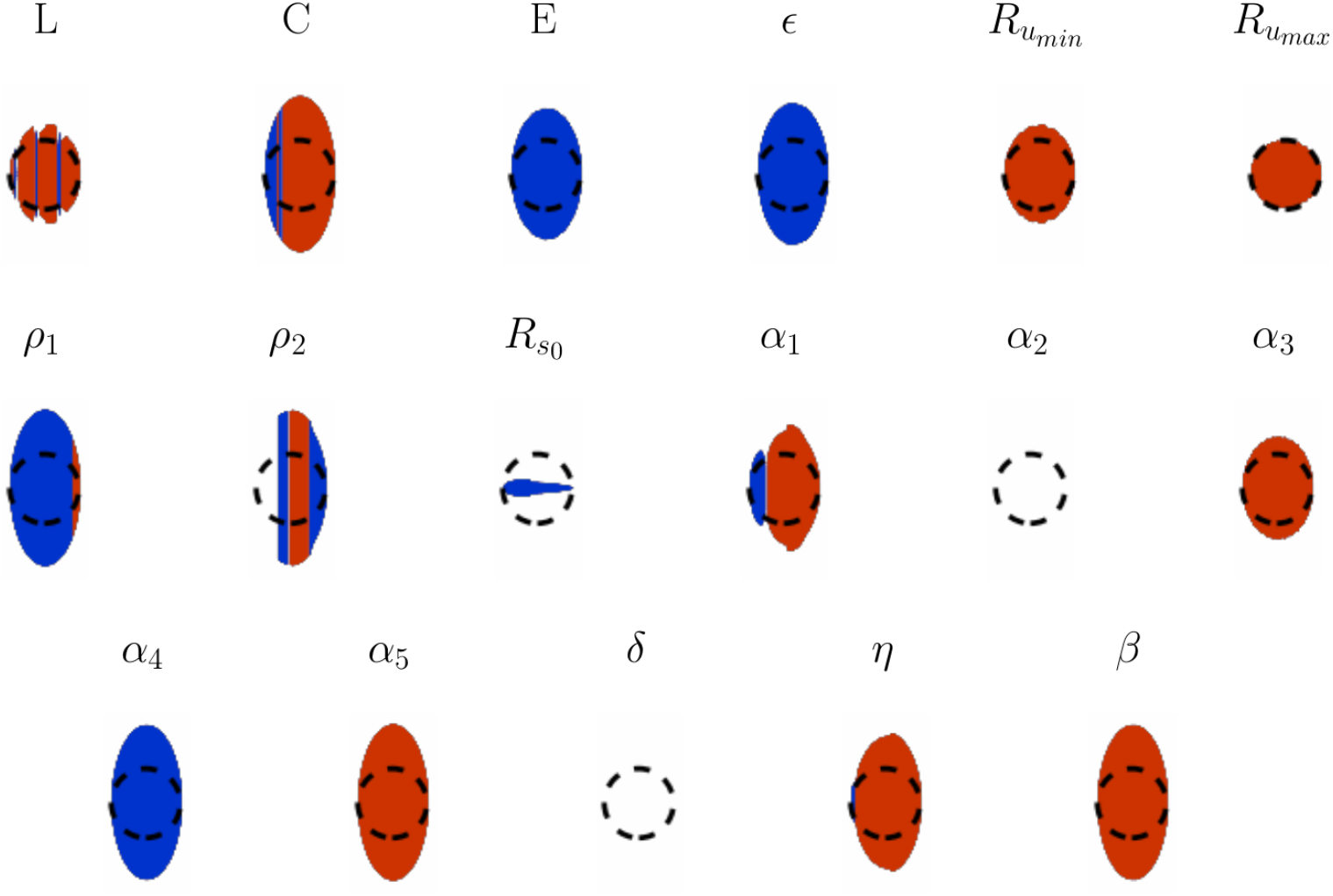
Parameters variation effect over the hard bump time. For each simulation, the rod has been exposed to a single *1μsec* flash causing the equivalent of 150 rhodopsin photoactivation. This diagram displays 2 non-impacting (*α*_2_ and *δ*), one low impacting (*Rs*_o_) and 14 strongly impacting xparameters. Two interesting phenomena are here represented. First, we observe the result of opposite parameter-value to effect-intensity relation between rise and decay times (see Figure 9 and 10) for *η, ρ*_1_ and *α*_1_ (bleu slice on the left hand side of the circle). Hence, even if only informative, this parameter-analysis technique seem to efficiently show a part of the topology of the parameter space. Second, the inductor (*L*) appears, when excited, to introduce noise in the system as its inductance value changes. This might be something to be careful about: in a network, the concurrent light activation and synaptic current feedbacks may amplify the latent instability.

**Figure 12.**
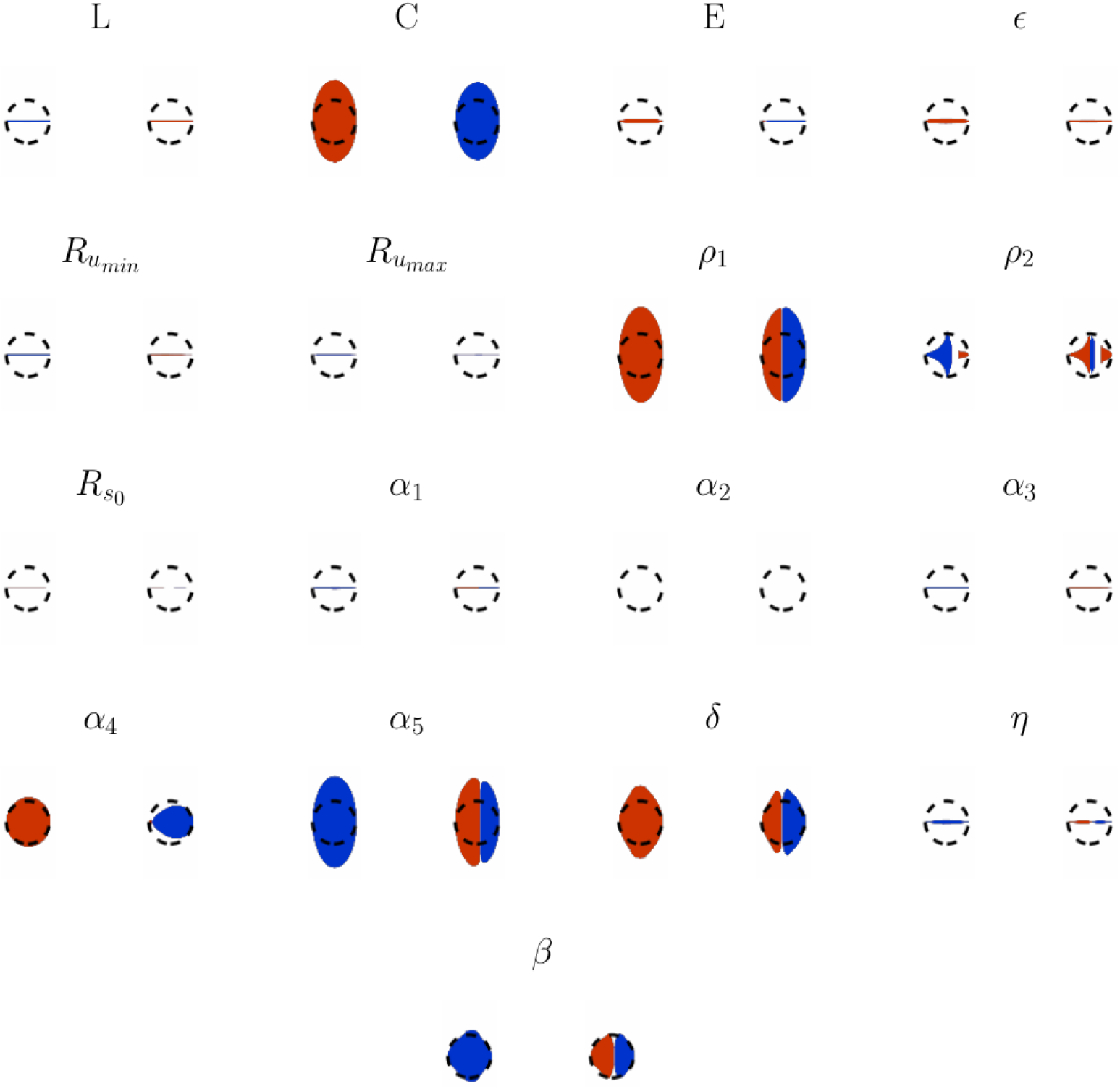
Parameters variation effect over the duration dependent response. For each simulation, the rod has been exposed to flashes of 1 to 500μsec long with an energy causing the equivalent of 150 rhodopsin photoactivation. The figure here displays for each variable the evolution of the average and standard deviation of the higher level of activation reached by the model over the different flash durations. Surprisingly enough, only six parameters (*C*; *rho*_1_,*α*_4_, *α*_5_, *δ* and *β*) display a high effect over the monitored feature. When taken in consideration with Figure 13, it appears that both effects can be decorrelated by modifying parameters *α*_4_ or *δ* as they present a stronger effect on the duration-dependent response than on the level-dependent activation.

**Figure 13.**
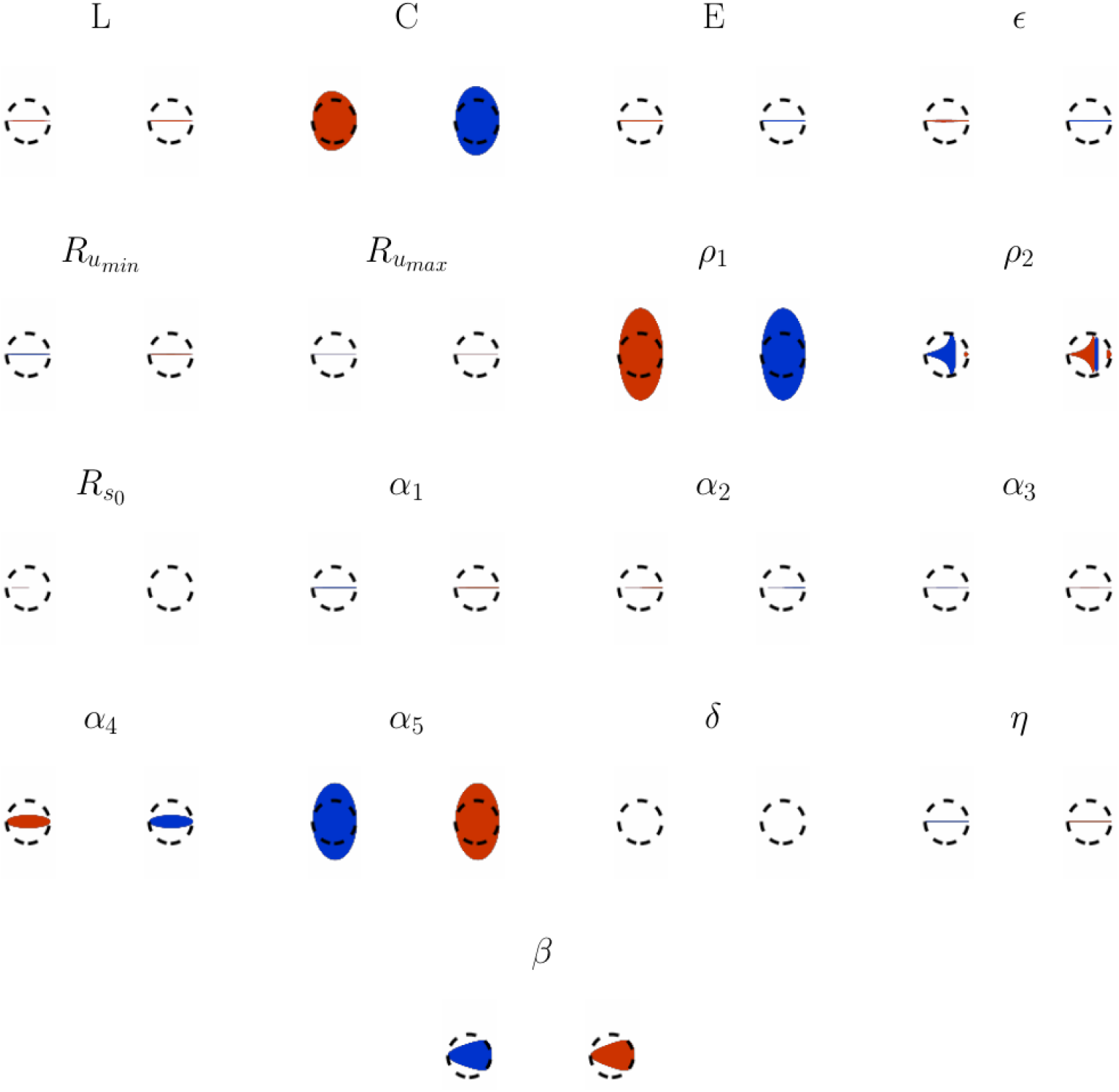
Parameters variation effect over the excitation level dependent response. For each simulation, the rod has been exposed to 1*μsec* flashes causing the equivalent of 1 to 1000 rhodopsin photo-activations. The figure here displays for each variable the evolution of the average and standard deviation of the higher level of activation reached by the model over the different flash intensities. Surprisingly enough, only four parameters (*C, ρ*_1_, *α*_5_ and *β*) display an high effect over the monitored feature. When taken in consideration Figure 12, it appears that both effects can be decorrelated by modifying parameters *α*_4_ or *δ* as they present a stronger effect on the duration-dependent response than on the level-dependent activation.

#### Rise, decay time and hard bump

Rise, decay time and hard bumps are features characteristic of the kinetic of the rod response due to a flash of light. As their name implies, we record the onset time of the cell response (see Figure 9) along with its time of decay (see Figure 10). However, in an attempt to obtain an indicator of both slopes we also record the time separating the half-rise level of the half-decay points (hard bump, see Figure 11). This indicator, even if mirroring non unique solutions for a single recording, is enough to give an insight on the shape of the curve assuming two constraints:

- The time to activation is always shorter than the recovery period, assuming the best set of parameters.
- The sampling of the parameter space is performed while making sure new parameters stay close enough from the best known set of parameters (see section *Methods*) to avoid heavy behavior alteration.

#### Duration, Excitation-dependent responses and damped oscillation

As previously mentioned, the response of our model is sensitive to both the strength of the input signal and its duration. Here, over a fixed signal intensity (150*γ* absorbed), we vary the duration to range between a single time quantum and 500 ms. Figure 12 presents the evolution of the average response level (left panel) along with its standard deviation (right panel). A similar method is applied to measure the intensity-dependent response level (see Figure 13). Here, many single millisecond signal events are sent ranging from 1 to 1000*γ* absorbed.

We use these mappings to represent the basic elements of a realistic light stimulation, including intensity and duration covariation, and study our model under punctual, yet continuous data flow.

Depending on the parameter values, a damped oscillation can also be visible in the early response, consecutive to any new steady excitation level. We, however, choose not to study this phenomenon in details here, as its origin can easily be imputed to the inductor of the RLC filter in the absence of feedback from the membrane model to the internal cell gear (ODE loop, see Equation 5). The ODE loop has indeed been designed to produce an hysteresis. The only element capable of expressing an oscillation appears to be equation 5. However, its activation level has been normalized on purpose (normalization term in Equation 5b) so that its early response cannot overshoot the steady state.

We have chosen parameter values for which any damped oscillation disappears after a demi-period helping the filter to create a solid bump of activity, in response to a flash excitation, but not allowing it to alter the full response shape.

#### Result representation

The studied behaviors suppose the general shape of the curve to be close enough to the response produced using the best known set of parameters. To ensure so, the range of sampling is chosen after the log scale of the best known parameters. Each of them vary over a range of values one log unit lower than its own log scale (see subsection *Effects comparison*). That way, we explore the parameter space neighbourhood of the best known values while remaining in a valid numerical space.

The measured effects are transformed (subsection *Capping*) and used as an area (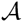 - see Equation 2) value for an ellipse of demi-major axis (or semi-minor axis, depending of the inferred size of its conjugate) 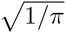 (the ray of a circle or unitary area). The horizontal axis stands for the space of parameter sampling and, as for, each point on this axis let appear its corresponding elliptical image for the computed 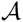.

The elliptical blobs of Figures 9 – 10, 11, 12 and 13 built after this method are concatenation of elliptic fragments representing the variation of the effect according to the values of a particular parameter projected on a circular/elliptical referential. Colors indicate a positive (red) or negative (blue) correlation between the variation of the effect and each parameter value. A sudden interruption of the blob reveals either an absence of effect variation or data (model divergence).

With that method, an effect linearly correlated to a parameter will display a regular ellipse; and, if their variations appear to be equal with respect to their initial values, a red circle of area 1 exactly fitting the dotted black circle present on each figure (see Figure 2).

#### Interpretation

By taking into consideration the effect on different parameters over the set of chosen features, it is possible to derive potential instability sources. Considering Figures 9, 10, 11, 12 and 13, it is clear that *ρ*_2_ is a parameter to carefully manipulate: As its value changes, the model appears to diverge with respect to hard bump time recordings. Additionally, the selected best known value seems to be an extremum when looking at the effect of this parameter over duration and level-dependent responses. As the value increases, the standard deviation of those effects quickly fluctuates suggesting a highly nonlinear parameter space.

Another potential instability point can be seen in the inductance value *L*. Along the range of alternative parameter values, its effect over the hard bump time appears to be non linear, and, when exposed to higher voltage or additional current sources (synaptic current), this phenomenon may spread towards higher values of the parameter and cause unseen aberration.

Another goal of our analysis was to give an insight into the topology of the parameter space. When looking at the parameter *α*_1_ over the rise and decay times, an opposite correlation can be identified. However, if the hard bump-time is taken in consideration, if appears that the negatively correlated effect visible on the rise time study is slightly stronger than its positively correlated component present of the decay time study; as a blue slice appears on the left hand side of the circle. This effect is also present on the same three experiments, in a noisier version, for the parameter *L*.

This method is of course perfectible, as the sample-range is solely determined from the log scale of the parameters best known value, without regard to its position within the log range. In other words, a value of 999 will be sampled as a second order of magnitude, whereas 1000 as a third order of magnitude, even though separated by a single unit. Additionally, we do not address the question of cross parameter variation when monitoring effects. Although of interest for an exhaustive description of behaviour, we chose to focus on the manipulation of single parameters and to reserve more complex manipulation for the future.

## Discussion

We described a qualitative model of the rod photoreceptor. Our modular representation includes a wide portion of the original cell behavioral panel and, unlike other models available in the literature, is able to perform when exposed to continuous input light signal. This aspect makes it fit to study the time-dependent mechanisms that can emerge within the retina when exposed to natural light streams.

With a view to studying retinal function, creating a model of rods is not useful on its own. One must be able to use it as a building block for a more complete representation of the visual processing stream. To do so, it is important to be able to modify expected behaviors of the model, and adapt the model to other needs [36], such as interacting with models of other types of cells, representing information transfer in damaged tissue, or in changing conditions, like room temperature and background luminance, etc.

This requirement, however, force modellers to favor fewer functional parameters over numerous biologically accurate variables. It is therefore all the more important to understand and predict behavior for the entire parameter set, over and beyond best known values.

This is the case for a lot of purely mathematical models (e.g., Fitzhugh-Nagumo model [43], Hindmarsh-Rose model [45]) as their parameters are abstract representation of chemical processes. To relax this constraint, we divided the model into three logical blocks (chemical light absorption, temporal loop, membrane) to facilitate the interpretation of results and manipulation of sets of parameters, whilst limiting systemic side effects. This deliberate choice is also representative of the fact that, even though the internal processes constitutive of the rod are relatively well understood, modelling a fully functional network of such cells is extremely costly [46].

One can also point out the absence of numerical accuracy in the model activation response. Biological neural networks are well known for their resilience, it has been also shown that for some networks, having a high enough number of components tends to fade out the irregularities between them (resistor, [47]). Hence, we hypothesize that having a super-accurate model might be irrelevant as the rods are, as any living cell in vivo, very different from each other. This might push the system towards a self-consistent referential based upon a limited set of features present within the neural response, rather than upon its absolute level of activation. Such models would not be very informative as artificially prescribed to a given situation.

The photoreceptive representation proposed by Clark et al. [33] is the closest model to what we propose. Their model is simple enough to allow formal mathematical analysis. In contrast, our model provides greater modularity, at the cost of mathematical simplicity, permitting the integration of additional mechanisms displayed by the real cell (slow dark adaptation, noise process, etc…). Our model focuses on the spread of electrical signal, and its interface with a network of cells is therefore as simple as it can be, based a proper membrane electrical representation.

In our case the coupling could be done by modifying the RLC filter as the synaptic current does not act on the phototransduction mechanism. A second variable generator could be added to the one present (E), standing for the current flowing though the particular gap junction synapse linking a photoreceptor to the rest of the retina in mammals [14].

Using the same idea, the system could be driven by two generator instead of one (a positive current source and a negative both gated by a respective resistor). Following Baylor, Nunn and Schnapf [39], by taking in account the Johnson-Nyquist noise in the positive gating resistor *R* using the Nyquist equation.

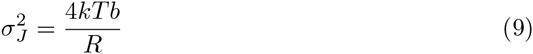

Where 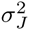 is the noise variance, *k* the Boltzmann constant, *T* the temperature in Kelvin and *b* the recording bandwidth. Upon light activation, the value of *R* would increase allowing the membrane to hyperpolarize (and consequently decrease the noise variance). This would be coherent with the phenomenon observed in vitro assuming dark adaptation: dark noise displays a high variance in the dark but decreases when the cell is stimulated [39]. This naturally leads to the question of light sensitivity. The quantum nature of light quantic interaction with the photopigment molecule induces a stochastic detection process. It this work, we positioned ourselves downstream to this process assuming that all absorbed photons went through the quantum induced uncertainty [5]. Further, the visual system itself appears to be a dynamic machinery able to modify its light detection capabilities according to the external light history. This process can be incorporated with an additional differential equation acting on the transition function from *v* into *R_s_* (See Equation 7). This equation would present a slower dynamic than displayed by the current ODE loop; of note, a total dark adaptation of the retina is typically achieved after 40 to 50 min [48].

Our model constitutes the first step towards a time-continuous, behavior-focused model of the retina. Our next step will be to create a representation of the horizontal cell layer aiming to interconnect artificial rods together before building higher retinal layers. At this stage, all our simulations have been ran at the scale of the micro-second, and, even-though the differential equations used here are fit to different time scales, we do not think that having a thinner temporal granularity would significantly improve the quality of the analysis but will surely increase the simulation time by a scale. A coarser time resolution, on the other end, would prevent consideration of the really brief flashes of light used in the empirical literature. The final model might thus be accurate and light enough to embark a reasonably large amount of neurons while enabling population activity study underlying early vision processes.

Our observation of this first layer of cells leads us to hypothesize that the first logical layer, comprising rods and horizontal cells, may have several functional roles. We observed a time and spatial high-pass filtering process consequent to both the horizontal cell negative feedback and the rod shape response. This first observation is, however, still to be studied. The band pass filtering property of our photoreceptor model could be reduced depending on the type of feedback and the sizes and shapes of receptive fields in the horizontal-cell layer. We expect this study to be the opportunity to refine our parameter sensitivity analysis by addressing the above mentioned limitations.

To conclude, our work stems from the hypothesis that there is more to the retina than the simple transduction of photonic energy, for transmission and processing by early visual areas of the cortex. Despite having been one of the most studied organs of perception, the retina is still primarily perceived as a gateway to the brain. Its computational powers are yet to be unravelled, and when empirical study may fall short, we hope that computational efforts will shed light on new avenues of research.

## Acknowledgments

This work was conducted as part of the doctoral thesis of TD, and funded thanks to University of Reading Research Award 2013–2017 to EBR. The funders had no role in study design, data collection and analysis, decision to publish, or preparation of the manuscript.

